# A novel averaging principle provides insights in the impact of intratumoral heterogeneity on tumor progression

**DOI:** 10.1101/584490

**Authors:** Marta Leocata, J. C. L. Alfonso, Nikos I. Kavallaris, Haralampos Hatzikirou

## Abstract

Typically stochastic differential equations (SDEs) involve an additive or multiplicative noise term. Here, we are interested in stochastic differential equations for which the white noise is non-linearly integrated in the corresponding evolution term, typically termed as random ordinary differential equations (RODEs). The classical averaging methods fail to treat such RODEs. Therefore, we introduce a novel averaging method appropriate to be applied on RODEs. To exemplify the importance of our method, we apply it in an important biomedical problem, i.e. the assessment of intratumoral heterogeneity impact on tumor dynamics. In particular, we model gliomas according to a well-known Go or Grow (GoG) model and tumor heterogeneity is modelled as a stochastic process. It has been shown that this GoG model exhibits an emerging Allee effect (bistability). We analytically and computationally show that the introduction of white noise, as a model of intratumoral heterogeneity, leads to a monostable tumor growth. This monostability behaviour is also derived even when spatial cell diffusion is taking into account.

## 1. Introduction

In the current section we demonstrate the key ideas of the novel averaging principle is introduced in the present work. For that purpose let us consider the following simple ordinary differential equation (ODE)

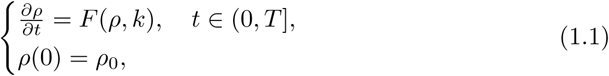

where *F*: ℝ × ℝ → ℝ is a globally Lipschitz function in both variables, *ρ, k* ∈ ℝ. Here the variable *k* represents a generic parameter of the model represented by the system (1.1). Assume now that the parameter *k* is not constant but varies randomly in time, and thus some noise is introduced to system (1.1). One can assume that the introduced noise resembles a Gaussian Process, or in more singular case, when it varies very strongly and at very short time scale, that has the form of a white noise. In the latter case and when the dependence on the parameter *k* = *k*(*t*) is linear, namely

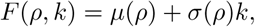

where then system (1.1) is reduced to the following initial value problem of a stochastic differential equation (SDE)

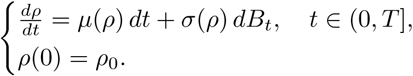

Next in the case the dependence on *k*(*t*) is non linear and *k*(*t*) is a real valued process, hence not white noise, then system (1.1) is well defined and its mathematical study can be delivered through a well established theory, see.^12^ Otherwise, if *k*(*t*) is thought as a white noise, namely as a uniform distribution will be denoted by *ξ*_*t*_ to make more clear the reference to a stochastic process, then we have to deal with the following system

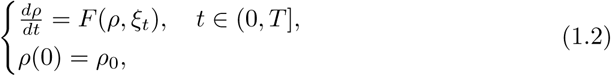

which is meaningless in the context of stochastic differential equations (SDES), since the drift of the ODE (1.2) is given by a non linear function of a distribution, namely of the white noise

To tackle system (1.2), the idea is to work with an approximation of the white noise *ξ*_*t*_, which is defined in a complete probability space (Ω, *𝓕*, (*𝓕*_*t*_)_*t*∈ [0,*T*]_, ℙ) with filtration (*𝓕*_*t*_)_*t*∈ [0,*T*]_. We will call one of the possible approximations of *ξ*_*t*_ as 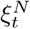. Then considering 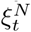, instead of *ξ*_*t*_, we obtain the following well posed system,

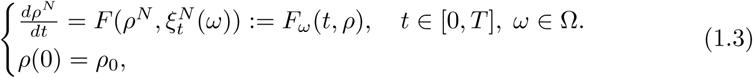

where the equation of (1.3) is a random ordinary differential equation (RODE), that is a non-autonomous ODE for almost every realization *ω* ∈ Ω.

The challenge is to investigate whether system (1.3) has a limit as *N* → ∞. To do so we will focus on a specific case, in particular we assume that

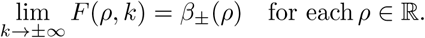

Intuitively speaking 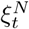 are independent centered Gaussian variables with variance that tends to infinite. Our main aim is to figure out how the nonlinear term *F* behaves when it is perturbed by an approximation of the white noise. Let us think the variable *ρ* being fixed, then the function *F* will be evaluated in extremely big positive values (on which *F* ≈ *β*_+_) or extremely big negative values (on which *F* ≈ *β*_−_).

Thus one would expect that in the limit *N* → ∞, *F* becomes a sort of Bernoullian process, denoted by *α*. Namely on a probability space (Ω, *𝓕*, (*𝓕*_*t*_)_*t*∈ [0,*T*]_, ℙ), the random variable *α* is defined by

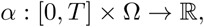

With

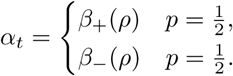

Furthermore for each *t*_1_, *t*_2_ ∈ [0, *T*], we expect 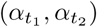 are independent random variables. Consequently, in the limit *N* → +∞ one would expect as a limit equation of (1.3) a RODE with a drift term which maintains a random oscillation between the two values *β*_+_ (*ρ*) and *β*_−_ (*ρ*). Surprisingly this is not the case and as we will see in section 4 the limit equation is a deterministic ODE, whose drift is given by the mean of the two extremes, i.e. 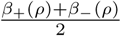.

The outline of the current manuscript is as follows. In section 2 for reader’s convenience we present some preliminary material. Section 3 introduces some auxiliary results, related with the averaging approach. The previous is followed for the demonstration and the proof of our main mathematical result, the novel averaging principle presented in section 4. In section 5 an application of the introduced averaging principle on the impact of intratumoral heterogeneity in glioma progression is provided. It is actually observed that the introduction of randomness in the intratumoral heterogeneity works towards the disappearance of the Allee effect. Finally we close the paper with a discussion of our main results in section 6.

## 2. Preliminaries

In the current section some preliminaries are presented for reader’s convenience. Throughout the manuscript we consider random approximations of a white noise (*ξ*_*t*_)_*t*∈[0,*T*]_. White Noise can have different equivalent definitions, however we will focus on a particular one: the one that allow us to figure out the white noise as the “derivative” of Brownian motion. Before proceeding with the required definitions we just point out that we will work in the filtered probability space (Ω, *𝓕*, (*𝓕*_*t*_)_*t*∈ [0,*T*]_, ℙ).

First, we recall the definition of Gaussian stochastic process generalized as can be found in.^7^

### Definition 2.1

Let us consider a generalized stochastic process Φ, namely

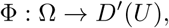

where *U* is an open set in ℝ and *D′*(*U)* is the space of distribution on *U*. We say that Φ is a gaussian process if for each 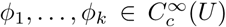 the random vector Φ*B*(*ϕ*_1_),…, Φ(*ϕ*_*k*_) is normally distributed.

Notice that a Gaussian process is characterized by the following quantities: its mean

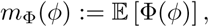

and its covariance

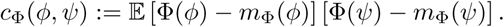

for each 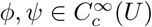.

### Definition 2.2

Let (*B*_*t*_)_*t* ∈ℝ^+^_ a Brownian motion on the probability space (Ω, *𝓕*, ℙ). Associated to *B* we can consider the generalized stochastic process,

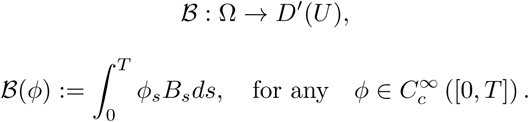

Then we define the white noise as the generalized derivative ℬ′: Ω → *D*′([0, *T*]) of the generalized Brownian motion 𝓑, that is

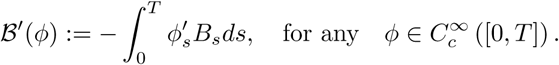

Henceforth we will denote 𝓕 ′ by *ξ* and thus

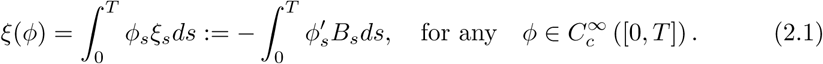

White noise is a generalized Gaussian Stochastic Process, since

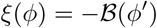

and 𝓕 is a Gaussian process. Recalling that cov(*B*_*t*_, *B*_*s*_) = min{*t, s*}, we can evaluate the mean and covariance of the white noise as

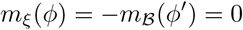

and

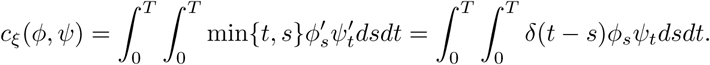

So we can conclude that the white noise is a *δ*-correlated stationary, Gaussian process with mean zero and covariance 𝔼 [*ξ*_*t*_ *ξ*_*s*_] = *δ*(*t - s*).

As it has been already mentioned at the beginning of the section our main strategy is to work with an approximation of (*ξ*_*t*_)_*t*∈[0,*T*]_, since only the associated ODEs are well defined.

Let us now consider the stochastic process 𝓕 ^Δ^, defined by the difference quotient of Brownian motion, 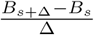, namely

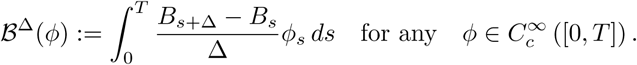

We justify our choice with the following lemma.

### Lemma 2.1

*The process* 𝓕 ^Δ^ *converges weakly to ξ, almost surely, that is*

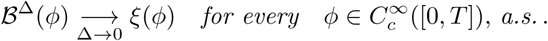

**Proof.** Due to the non-differentiability of Brownian motion we first prove the lemma for a mollified approximation of Brownian motion 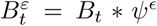, where ψ^*ε*^ is a mollifier function, see,^15^ and then taking the limit as *ε* → 0 we will deduce the desired result. It is easily seen that for any fixed;ε we derive

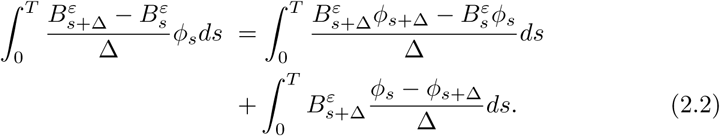

Taking now the limit of (2.1) as Δ → 0 we derive

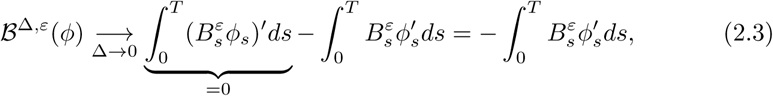

since 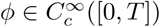.

Finally considering the limit of (2.3) as *ε* → 0 we derive the desired result.

Now we consider a discrete sampling of the process 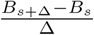 with Δ = Δ_*N*_ =1/N Thus we divide the interval [0, *T*] in small sub-intervals such that 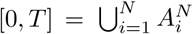 where 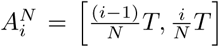 with *i* = 1,…, *N*. Note that in the interval 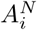 the sample is normally distributed

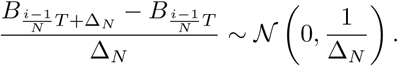

Consequently, inspired by inspired by Lemma 2.1, we consider the following random step function as an approximation of the white noise

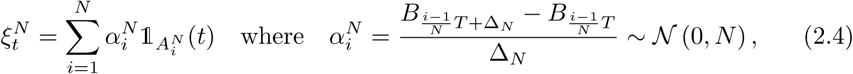

see Fig. 1.

**Fig. 1:**
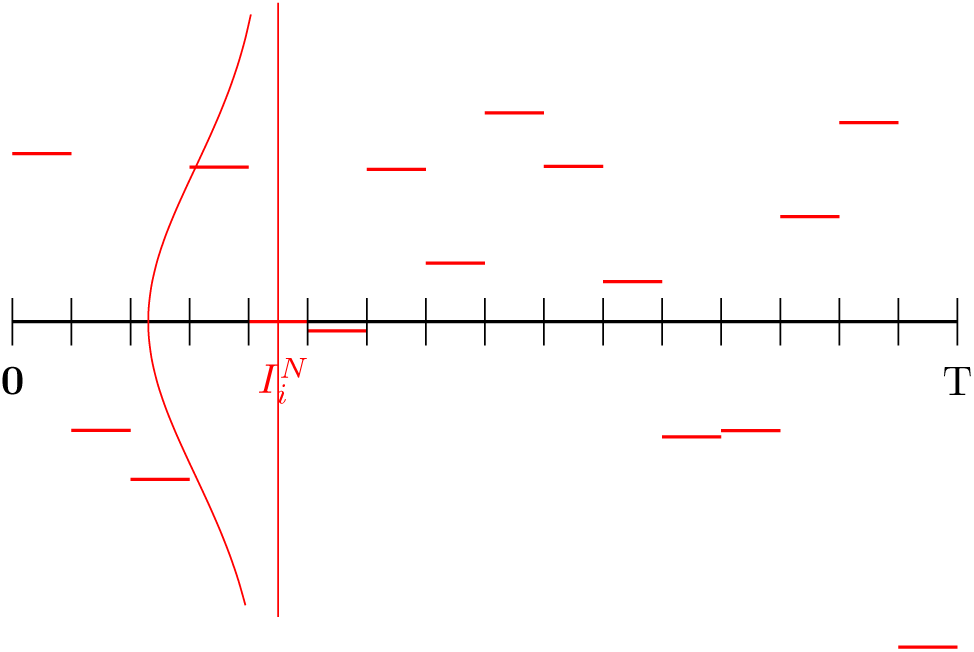
Plot of the approximation 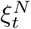 of white noise *ξ*_*t*_.

## 3. Some auxiliary results

Our mathematical approach to study system (1.2) is inspired by the averaging theory. Indeed, this theory treats dynamical systems in which two variables (*X*_*t*_, *ξ*_*t*_) coexist and interact, but in two different time scales. In particular, the time scaling of one of the two variables, *ξ*, is subjected to acceleration by a factor *ϵ*, which implies that its actual time scale is *t/ϵ* and so it contributes to the dynamics of the system in the form *ξ*_*t/ϵ*_. Via averaging theory we can investigate what is the impact to the system for small *ϵ*, namely when the acceleration is big, and we can conclude that the contribution of the faster process *ξ*_*t/ϵ*_ is evaluated in its time average.

Throughout the current manuscript we will try to implement the mentioned ideas into system (1.2). Since the latter system is not well defined when the random perturbation is a white noise we will deal with the approximating system (1.3), where the white noise is approximated by 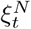, defined by (2.4), and the parameter of approximation *N*, can be reinterpreted as the new scaling time parameter^4^. In the current section we present some auxiliary results will be used for the proof of our novel averaging principle demonstrated and proven in section 4.

We now present the main hypotheses regarding the drift *F* term. Please note that *C* denotes a positive constant independent of *t* which might change its value from line to line.

### Hypothesis 3.1

1. *F*: ℝ × ℝ → ℝ is Lipschitz with respect the first variable, uniformly on the second, namely there holds

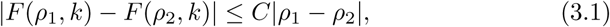

for each *k* ∈ ℝ, where the positive constant *C*, does not depends on *k*.
2. *F* is also bounded, i.e.

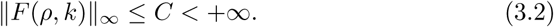
3. *F* is quasimonote, i.e.

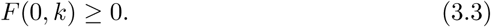
4. Finally for each *ρ* ∈ ℝ the asymptotic profile of the drift term is defined as

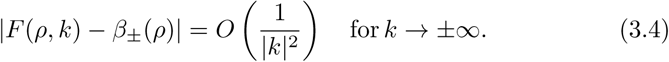

The way we interpret the solutions of RODE (1.3) as well as some of its features are provided below:

### Definition 3.1

By a solution of the RODE (1.3) we mean a stochastic process such that for each *ω* ∈ Ω, *ρ*^*N*^ (*ω*) ∈ *C*([0, *T*]) and satisfies:

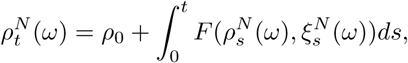

with a deterministic initial condition *ρ*_0_ ∈ ℝ^+^.

### Remark 3.1

Under the assumption that *F* locally Lipschitz in its first variable then problem (1.3) has a unique pathwise solution *ρ*^*N*^ (*t, ω*) for every *ω* ∈ Ω, in the finite interval [0, *T*],.^12^ Moreover the assumption (3.3) guarantee the positivity of the solutions of (1.3) initiated from positive intial condition *ρ*_0_ *≥* 0, see also Theorem 2.4 in,^12^ which is a desired property in biological models like the one that will be considered in section 5.

In the next section, see in particular Theorem 4.1, we prove that the limit towards *N* → +∞ of problem (1.3) is the following deterministic problem

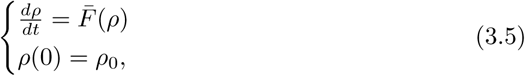

where

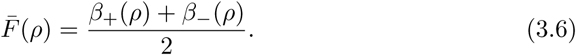

### Definition 3.2

By a solution of the ODE (3.5) we mean a function *ρ* ∈ *C*([0, *T*]) satisfying:

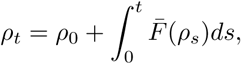

with a deterministic initial condition *ρ*_0_ ∈ ℝ^+^.

Again if the drift term 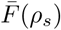 is locally Lipschitz continuous and positive, then problem (3.6) has a unique positive solution.

The rest of this section is devoted to the study of the average in time contribution of the process 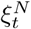 on the drift *F* (asymptotically in *N*), more precisely see Proposition 3.1. Before proceeding with the proof of this result we need a key tool, i.e. a law of large numbers for random variables (r.v.) 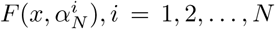, which is stated and proven in the following lines.

### Lemma 3.1

*Assume that F satisfies Hypothesis 3*.*1*. *Fixed N* ∈ ℕ, *we consider* 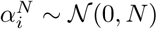 *as independent random variables, then for each x* ∈ ℝ^+^ *there holds*

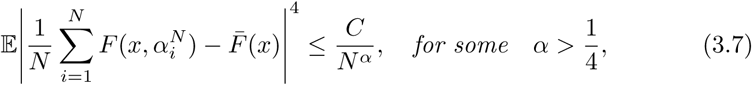

*where* 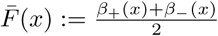 *and provided that N is sufficiently large*.

**Proof.** Let us call 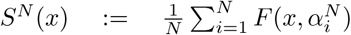, then for 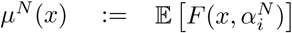, *i* = 1, 2, …, *N*, we have

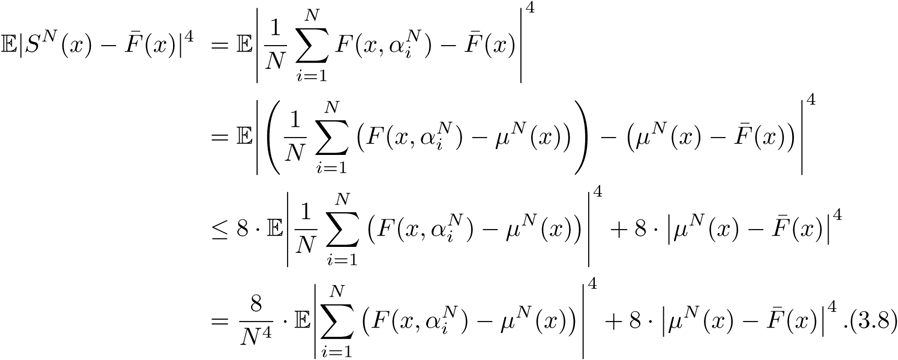

Now, setting 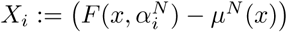 then the first term in the r.h.s of (3.8) can be expanded as

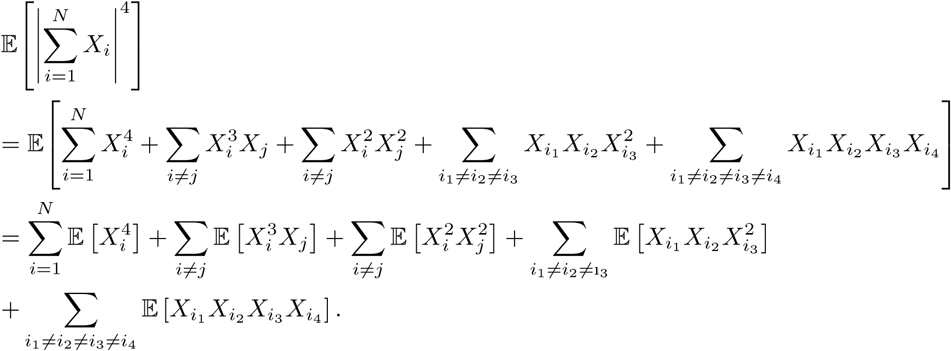

Due to the independence of the involved r.v. we derive

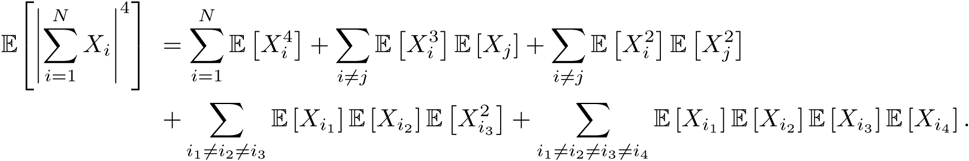

Since 𝔼 [*X*_*i*_] = 0, the second, the fourth sum and the last sum are equal to zero. So the previous quantity is reduced to

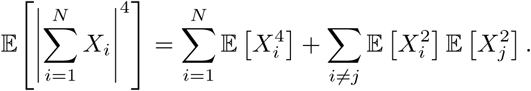

Thanks to the previous computation and to the fact that the r.v. *X*_*i*_ are independent and identically distributed (i.i.d.) and uniformly bounded with respect to *N*, we obtain:

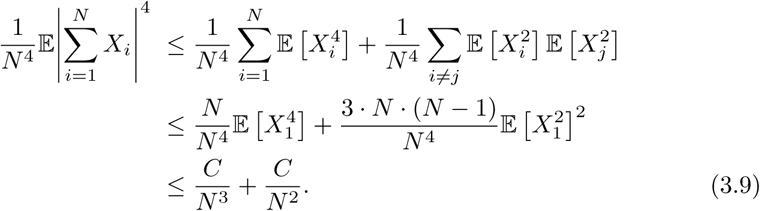

Next we try to estimate the second term in the right hand side (r.h.s.) of (3.8). It remains to prove that

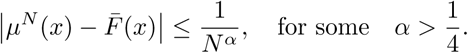

In the following it will be more convenient to rewrite 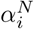 as 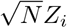, where *Z*_*i*_ ∼ 𝒩 (0,1) for *i* = 1, 2,*…*, *N*. Then considering the sequence *ϵ*_*N*_ = *N* ^*-β*^ with 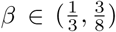 we have

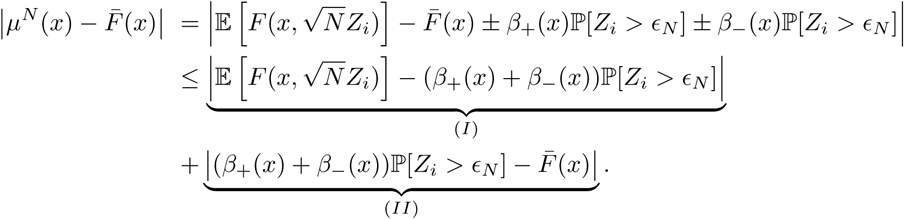

The first term (*I*) is estimated as follows

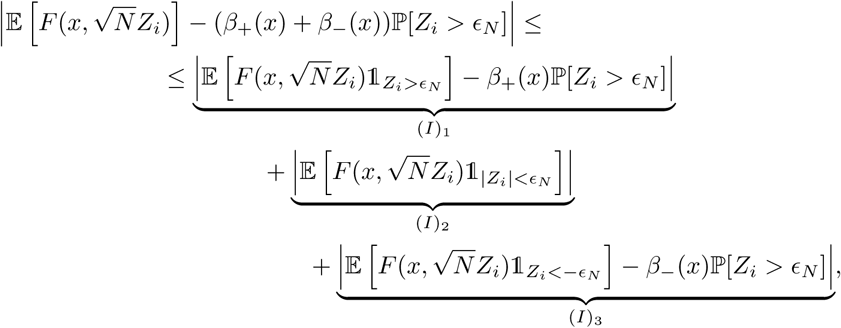

where (*I*)_1_ can be expressed as

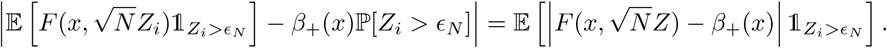

Due to condition (3.4), there exists 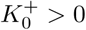 such that for each 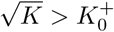

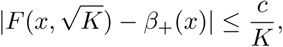

thus choosing *N* such that 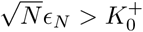, we have

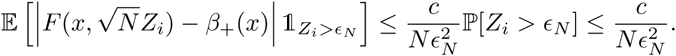

Recalling that *ϵ*_*N*_ = *N* ^*-β*^ we have that 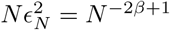 and thus

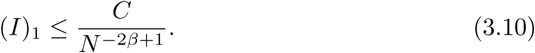

We proceed in a very similar way to estimate (*I*)_3_, which again can be written as

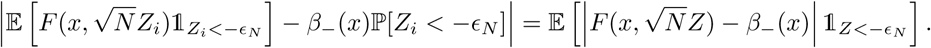

Again condition (3.4) entails the existence of some 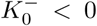 such that for each 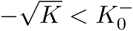 there holds

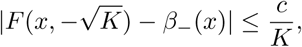

and thus by choosing *N* such that 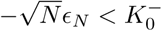, we obtain

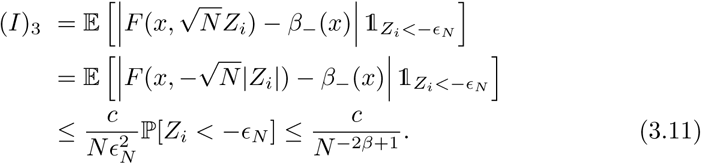

Regarding the estimation of term (*I*)_2_, since 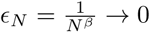 as *N* → ∞ and ℙ [*Z*_*i*_ = 0] = 0 then we can find *N*_1_ such that for *N* > *N*_1_

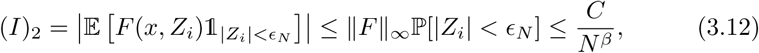

taking also into account (3.2). Regarding the last estimation, about term (*II*):

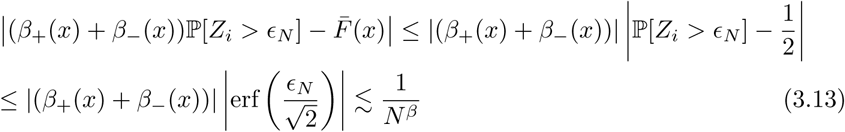

In summary, by virtue of (3.10), (3.11), (3.12), (3.13) and for 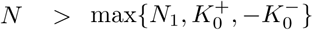 we deduce

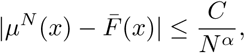

where *α* = min{*β*, 1 − 2*β*} = 1 − 2*β* > 1*/*4 for 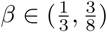. Thus in conjunction with (3.9) we finally obtain the desired relation (3.7).

Now we are ready to prove that the sequence of solutions *ρ*^*N*^ of system (1.3), converges to the solution *ρ* of system (3.5) whose drift is given by Definition 3.2.

### Proposition 3.1

*Assume that F satisfies Hypothesis 3*.*1*. *Let* 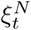 *be the approximation of white noise, defined in* (2.4) *then for each t*, Τ > 0 *and x* ∈ ℝ *there holds*

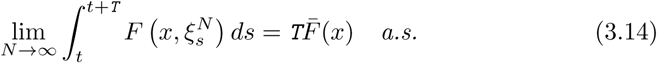

*with* 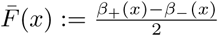.

**Proof.** Note that by the definition (2.4) of the process 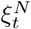 we immediately have

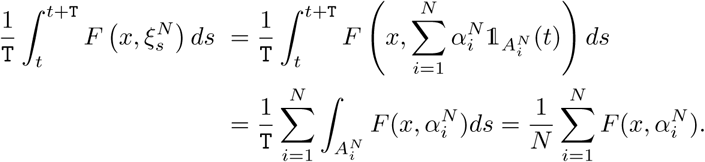

Next we prove that the sum 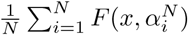 converges almost surely to 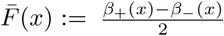. To this end we can not directly apply the classical result of law of large numbers, since we are treating a triangular sequence of random variables; namely notice that r.v. 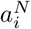 depend on the double index (*i, N*). In the following, for simplicity we will denote with 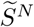 the quantity, whose limit is under investigation,

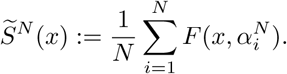

In order prove that 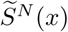 converges almost surely to 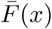 we equivalently demonstrate that there exists an infinitesimal sequence *ϵ* _*N*_, such that

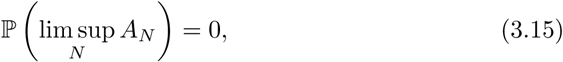

with 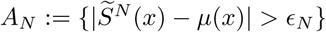. To this end we will use Borel-Cantelli Lemma and hence it is sufficient to show the convergence of the series 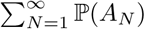. Next by virtue of Markov inequality we derive

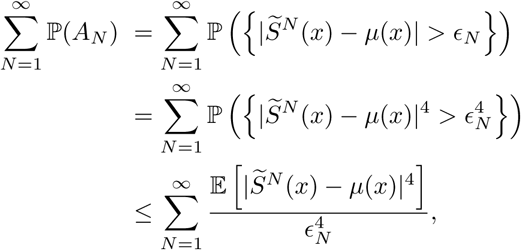

and thus by Lemma 3.1 we obtain

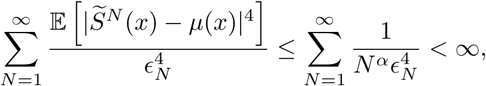

for some 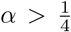. Then under the choice 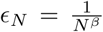 and imposing 4*β < α* − 1 we guarantee the convergence of the series. As a consequence we derive the validity of (3.15) and thus the proof is complete.

## 4. A novel averaging principle

In the current section we present and prove our main mathematical result. Indeed, using the auxiliary results provided in section 3, we demonstrate a novel averaging principle can then be applied to the approximation (1.3) of system (1.2). As it has been already explained in the introduction the investigation of the dynamics of nonlinear system (1.2), under a white noise perturbation, passes through the study of the dynamics of its approximation (1.3) where the white noise *ξ*_*t*_ is substitute by the sequence 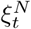 defined by (2.4). Then this idea can be applied, see section 5, to study the dynamics of some nonlinear models describing the tumor growth of a brain tumor (glioblastoma).

Our main mathematical result is stated as follows.

### Theorem 4.1

*Let* 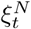 *be the approximation of white noise, given by* (2.4). *Then the solution trajectories* 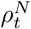 *of the RODE* (1.3) *converge uniformly in time and almost surely to the solution trajectories of the ODE* (3.5), *i*.*e*.

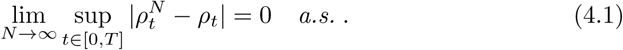

**Proof.** Let us take *ω* ∈ Ω. By the integral formulation of (1.3) and (3.5) we have

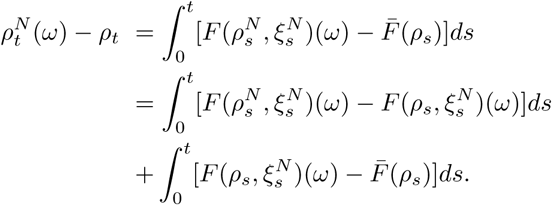

By considering the Lipschitz property (3.1) of the drift term *F* with respect to the variable *ρ* we obtain

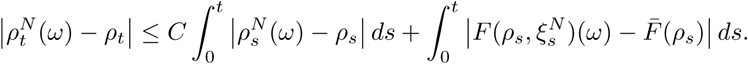

Applying now Gronwall’s lemma, see, ^14^ on the quantity 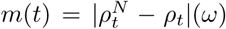 we derive

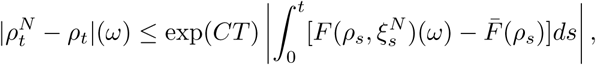

and thus

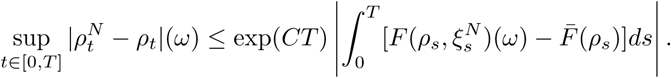

Hence in order to prove (4.1) it is sufficient to show that

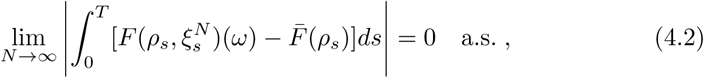

or equivalently to demonstrate that for any fixed *ω* ∈ Ω and for each *ϵ* > 0 there exists 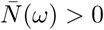, such that for any 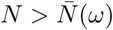 there holds

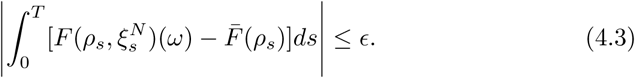

Now we fix *ϵ* and we discretize the interval [0, *T*] in *n* subintervals of amplitude 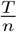, so we have

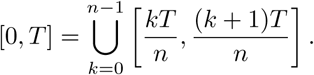

We underline the fact that the choice of *n* depends on *ϵ*, that is as small is *ϵ* then as big we must choose *n*. Using now the introduced discretization we have

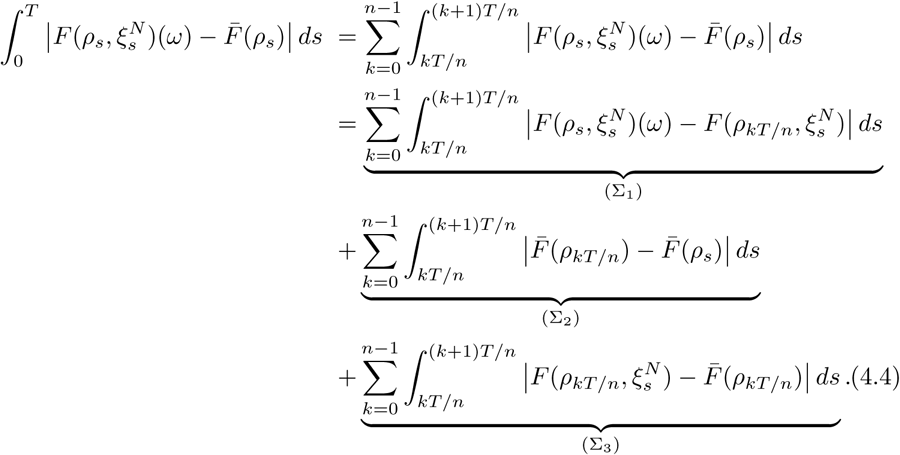

We first deal with terms (Σ_1_) and (Σ_2_) into relation (4.4). Due to the Lipschitz property of 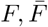 and *ρ* we obtain

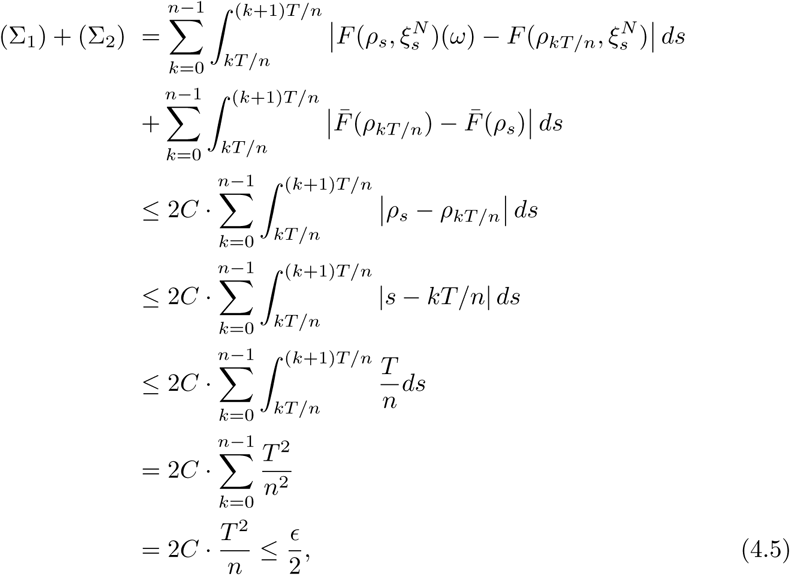

provided that 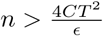, where *C* is a positive constant associated with the Lipschitz constants of 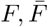 and *ρ*.

Regarding now the estimation of term (Σ_3_), we plan to apply Proposition 3.1 on each

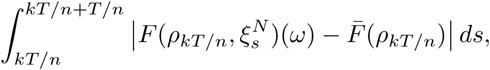

for *k* = 0,…, *n* − 1.

We choose to work on a subspace of measure 1, indeed we consider 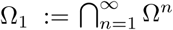 with 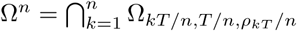 and where Ω_*kT/n,T/n,ρkT*_ */n* is the subsetof Ω on which the convergence result (3.14) holds for *x* = *ρ*_*kT/n*_, *t* = *kT/n* andΤ = *T′n*.

Now (3.14) entails that for any fixed *ω* ∈ Ω_1_ and for each 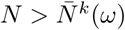

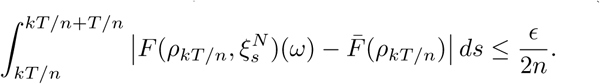

Consequently, by choosing 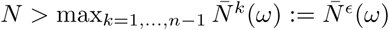 we deduce

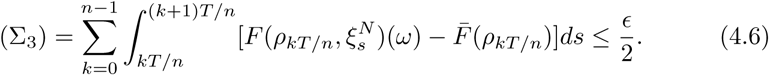

In summary, combining (4.5) and (4.6) then for any fixed *ω* ∈ Ω_1_ and for each *ϵ* > 0 there exists 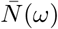 such that (4.3) holds for any 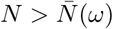

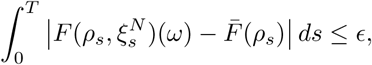

and thus the proof is complete.

Next we focus on the infinite dimensional case, i.e. we consider the following partial differential equation (PDE) problem

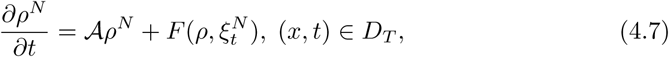

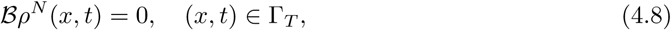

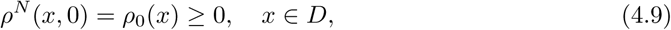

where *D*_*T*_ := *D* × (0, *T)*, Γ_*T*_ := *∂D* × (0, *T*) is a bounded and smooth *D* ⊂ ℝ ^𝓁^, 𝓁≥ 1 and *T* > 0 stands for the maximum existence time of solution *ρ*^*N*^, which is positive in *D*_*T*_ as long as it exists. Here 𝓐 represents a bounded operator who generates a strongly continuous semigroup *S*_𝓐_ (*t*) whilst 𝓑 is a bounded boundary operator. Besides the nonlinearity 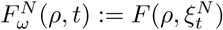 is assumed to be almost periodic with respect to *t* uniformly for *ρ* in compact subsets in Banach space *X*.

Consider now the averaged (PDE) problem

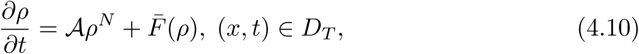

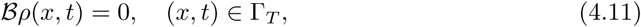

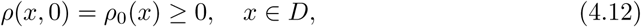

where 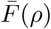 is given by (3.6). By virtue of a semigroup approach we can easily obtain a unique, positive solution for (4.10)-(4.12), see.^14^

Now by using Theorem 4.1 in conjunction with the semigroup approach introduced in^10^ we deduce the following infinite dimensional averaging principle

### Theorem 4.2

*The solution trajectories* 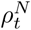 *of problem* (4.7)*-*(4.9) *converge uniformly in time and almost surely to the unique solution of problem* (4.10)*-*(4.12).

A special version of Theorem 4.2 is proven, by using an compactness approach, in section 5 see Theorem 5.1.

## 5. Application: impact of intratumoral heterogeneity in glioma progression

In this section, the main purpose is to use the averaging principle, as introduced in the previous section by Theorems 4.1 and 4.2, to address the problem of intrinsic heterogeneity in glioma progression. Gliomas originate from glial cells in the brain and constitute the most common type of malignant brain tumor in adults.^9^, ^23^ The WHO (World Health Organization) classifies gliomas into different classes, ranging from low-grade to high-grade malignancies^16^ depending on their proliferative potential and invasiveness. In particular, high grade gliomas are commonly characterized by diffusive infiltration into the brain tissue, behavior associated with poor prognosis and limited treatment outcomes. Glioma treatments usually involve resection accompanied by radiotherapy and′or chemotherapy. Herein, we use a mathematical model to understand the reasons of treatment failure and optimize the combination of current treatment modalities.

Two of the most important reasons for glioma treatment failure is due to intratumoral heterogeneity comprised by intrinsic heterogeneity and phenotypic plasticity.^17^, ^18^ The former describes the typically irreversible genetic or epigenetic diversity of tumor cells (e.g. due to mutations or clonal selection), and the latter is related to the reversible phenotypic responses towards microenvironmental cues. In gliomas, the most predominant phenotypic plasticity phenomenon is related to the “Go or Grow” (GoG) mechanism or migration′proliferation plasticity.^13^ This mechanism implies a mutually exclusive switching between migratory and proliferative phenotypes. The key question is how this tumor cell decision-making mechanism is regulated, and what is its impact on glioma growth and invasion. Concerning regulation, it was discovered a dependence of this cell decision mechanism on the local cell density without concluding on the exact functional form, by analysing images of *in vitro* experiments.^25^ Analysing further the potential local cell density dependencies, it was also found out low-grade tumor micro-ecology potentially exhibits an emergent Allee effect, i.e. a critical tumor cell density implying both tumor growth and control.^2^ The precise quantification of this critical tumor cell density could be a relevant prognostic criterion for the tumor fate, since it can be measured in biopsies samples. Moreover, it has been shown that this GoG mechanism explains the fast tumor recurrence time of high-grade brain tumors after resection.^13^ Finally based on our theoretical understanding of the cellular mechanism, it has been shown how personalized vasomodulatory cancer therapies can be optimized.^6^, ^11^ In particular, it has been revealed that one-size-fits-all vaso-modulatory interventions should be expected to fail, because control of glioma invasion characteristics, such as tumor front speed and infiltration width, can be very variable and may require more personalized therapeutic interventions.^8^

All the aforementioned models assume that all glioma cells have an identical GoG mechanism. However, in reality, each cell may have an idiosyncratic migration and proliferation regulation, following the GoG mechanism, due to intrinsic heterogeneity. The question is how we can model and analyze the impact of intrinsic heterogeneity of a tumor cell population, when migration and proliferation are regulated by a non-uniform GoG mechanism. In particular, we will focus on the existence of the Allee effect in the presence of intratumoral heterogeneity.

### 5.1 The Go or Grow model of glioma

For the reader’s convenience, we briefly present the derivation of the Go or Grow model, as introduced in.^13^ Due to the migration / proliferation dichotomy, we can distinguish the total population of glioma cells *ρ* in two groups, *ρ*^*p*^ proliferating cells and *ρ*^*m*^ migratory cells. For each type of population we can then write down an equation describing the corresponding dynamics, so we have the following system, which is described as a GoG model,

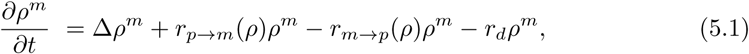

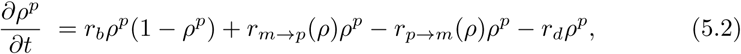

where *r*_*b*_ is the birth rate, *r*_*d*_ is the death rate, *r*_*p→m*_ (*ρ*) is the switch rate from proliferating to motile phenotype and *r*_*m→p*_ (*ρ*) is the switch rate from motile to proliferating phenotype of the tumor cells.

To obtain a unique equation for 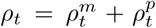, we refer to detailed balance condition

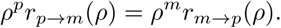

Through this condition we can rewrite the preceding system as a single PDE

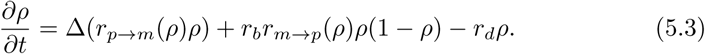

The phenotypic switching is regulated by local microenvironmental cues lumped into local density dependence. For the sake of simplicity, we will assume that both rates depend monotonously on cell density, approximated by a sigmoidal function with slope given by *k*. Intuitively, the slope can be viewed as the way that single tumor cell interprets its microenvironment and decides over its phenotype. Following,^2^ we consider that the two rates are complementary, namely if cell motility increases with cell density then cell proliferation decreases with density and vice versa: namely if *k* denotes the slope of the switch from motile type to proliferating type, then *−k* denotes the slope of the switch from proliferating type to motile type. A possible choice is given by the following, see Fig. 2:

**Fig. 2:**
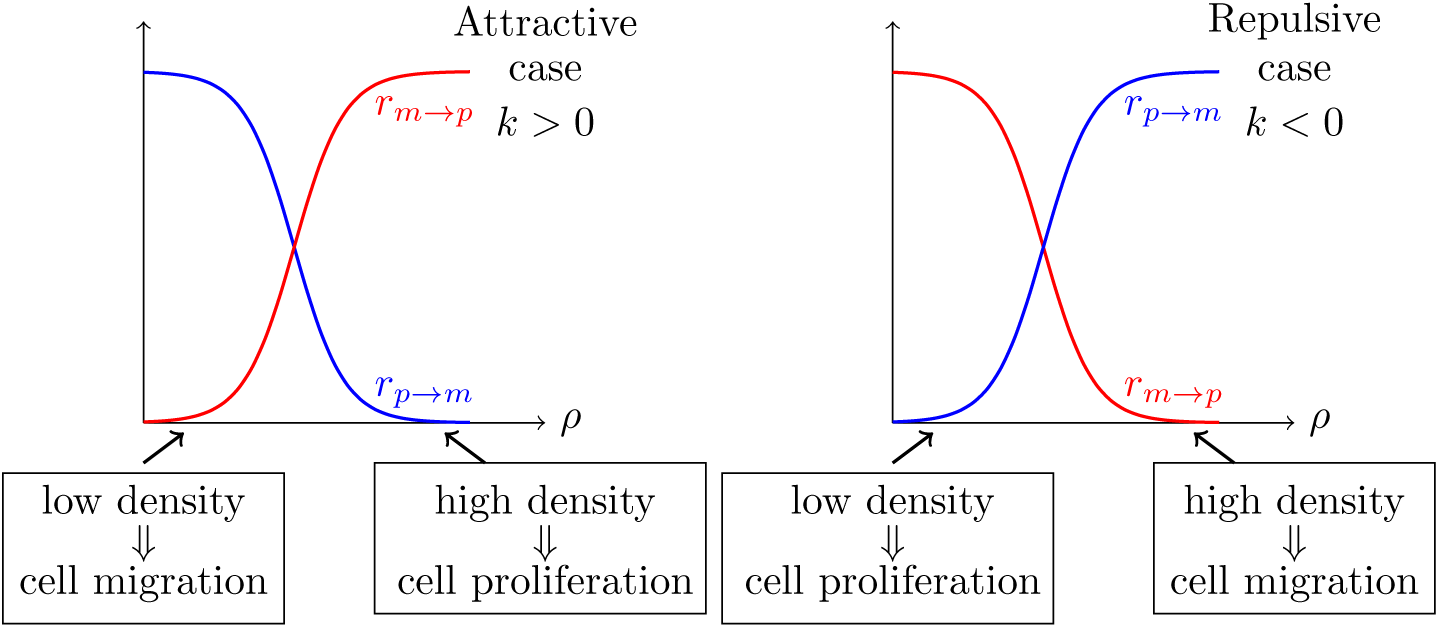
Sketch of cellular mechanism for different kind of phenotypic plasticity. In the left aggregative configuration is represented, while in the right figure, the repulsive configuration is represented.

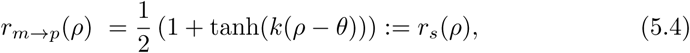

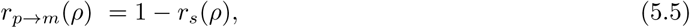

and thus (5.1) takes the form

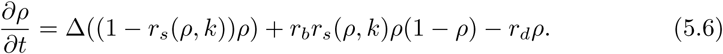

For positive slope *k* > 0 the phenotypic switch presents an attractive behavior, while for *k <* 0 a repulsive one (see Fig. 2).^2^ To the best of our knowledge the GoG model (5.6) has been only investigated for the case when *k* is a constant, see,^2^ i.e. the tumor cell population decides in a homogeneous way over proliferation or migration. Here we assume that tumor is heterogeneous in the way cells regulate their migration′proliferation phenotype controlled by a stochastic *k*.

### 5.2 Intrinsic intratumoral heterogeneity of glioma cells as a white noise

We recall that the sign of *k* indicates the regime where we are, it identifies if we are in an attractive or repulsive regime, whilst the absolute value of *k* measures the intensity of the phenotypic switching. In the following, we introduce the desirable heterogeneous regulation of the switch by assuming that *k* follows a probability distribution, i.e. we heuristically take

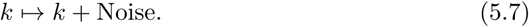

It is anticipated that the introduced intrinsic heterogeneity facilitates the tumor growth and persistence.^17, 18^ Therefore it is plausible to consider the “worst” case heterogeneity scenario, hence *k* is considered as a white noise. As first step towards the investigation of the dynamics of the GoG model (5.6) under random perturbation (5.7), we choose to neglect the contribution of the diffusion, and thus we initially consider the following ODE

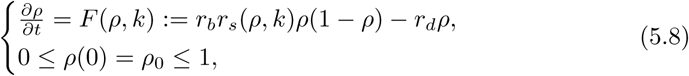

where

*r*_*s*_ (*ρ, k*) given by (5.4)

for *r*_*b*_ > *r*_*d*_, *θ* ∈ [0, 1] and *ρ*_0_ ∈ ℝ^+^.

If one wants to approximate *k* as a white noise, clearly relapses in the case exposed in the introduction, since the occurring system

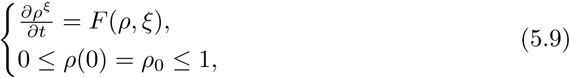

is not well defined.

In order to tackle system (5.9) we appeal to the averaging principle demonstrated by Theorem 4.1. To this end we just need to verify all the involved conditions of Hypothesis 3.1. We first note that condition (3.1) is trivially verified, since the drift term *F* (*ρ, k*) = *r*_*b*_ *r*_*s*_ (*ρ, k*)*ρ*(1 − *ρ*) − *r*_*d*_ *ρ* for *ρ* ∈ [0, 1] is differentiable. Besides *F* is bounded on both variables (*ρ, k*), since *k* appears as an argument of the hyperbolic tangent, and the variable *ρ* varies in a compact set. Thus condition (3.2) also holds. Obviously we have *F* (0, *k*) = 0 and thus condition (3.3) is also fulfilled. It remains to check the validity of condition (3.4). It can be easily seen that

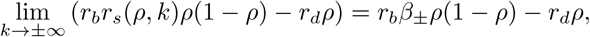

with *β*_+_ = 1 and *β*_*-*_ = 0 where the order of convergence is exponential, and hence condition (3.4) is also fulfilled. Consequently we have the following result which is a straightforward consequence of Theorem 4.1

#### Proposition 5.1

*Let* 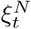 *be the process described in* (2.4). *Then the solution trajectories* 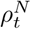 *of*

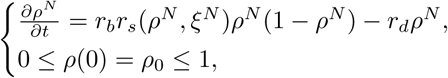

*converge uniformly in time and almost surely to the solution ρ of the following ODE problem*

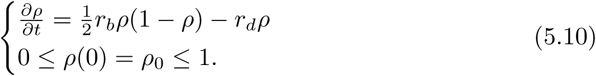

*Namely*,

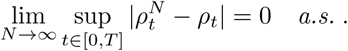

Thus in order to investigate the model (5.8), Proposition 5.1 provide us with the appropriate approximating ODE (5.10) which will be analyzed with respect to tumor steady states.

Let us here recall that in the absence of intrinsic heterogeneity, the trivial solution *ρ*_*nc*_ = 0 is a stable point for a certain range of parameters, as shown in Fig. 3 and.^2^ However, under the assumption of white noise intrinsic heterogeneity, the stability analysis of (5.10) leads to a monostable configuration. Specifically, the point 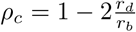 is the only stable point, whilst *ρ*_*nc*_ = 0 is still a fixed point but unstable (see Fig. 4). In this case the parameters (*k, θ*) do not influence the model’s steady states as in the deterministic one (see Fig. 3). The strength of the white noise is so large, that annihilates the existence of multiple steady states. Finally, the Allee effect that was observed in the deterministic case^2^ now disappears and the only remaining effect is the survival of a steady cancerous population.

**Fig. 3:**
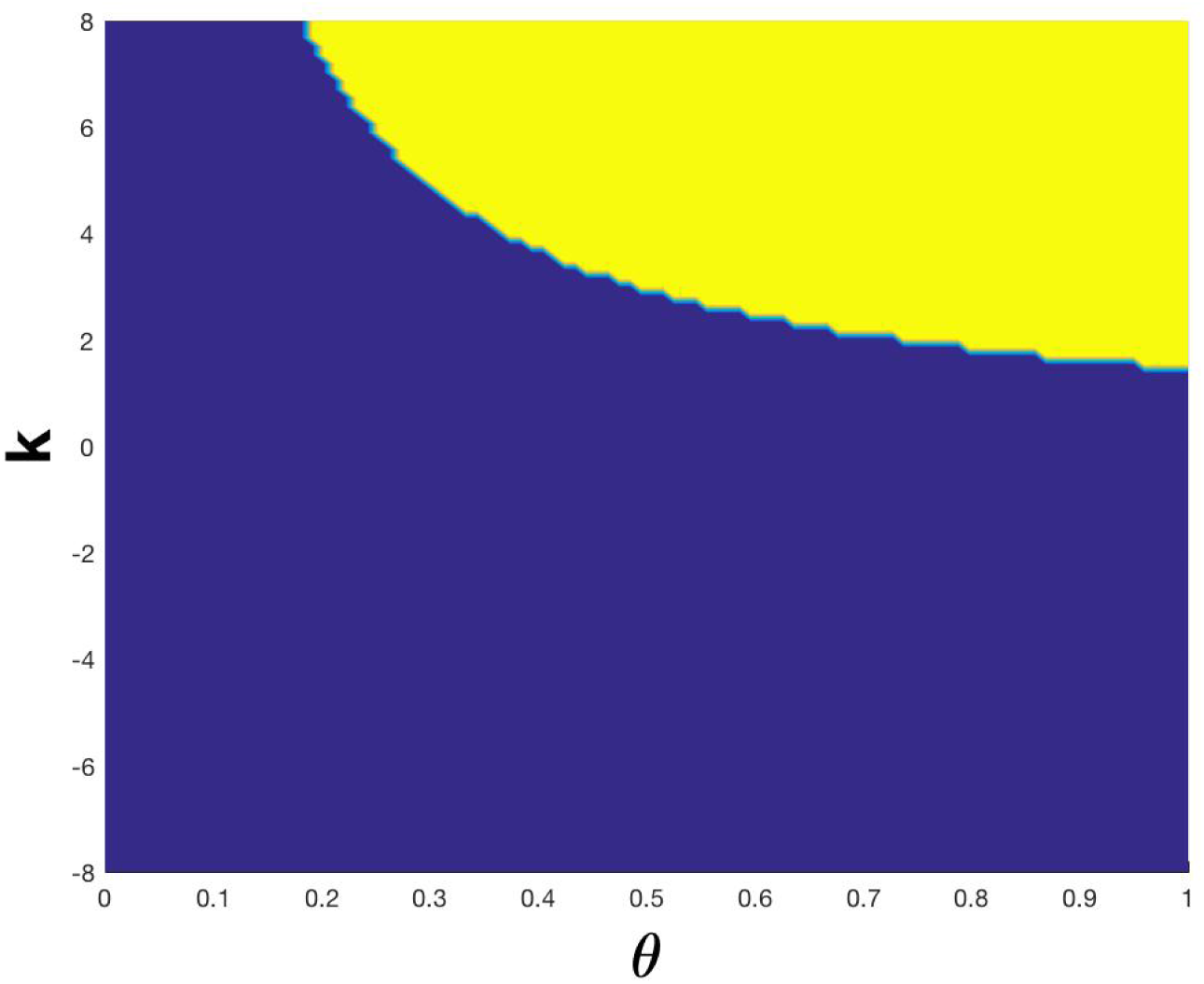
Allee effect in the deterministic system: The yellow area represents the area where 0 is stable for the above system, whereas the blue one depicts the area where 0 becomes unstable.

**Fig. 4:**
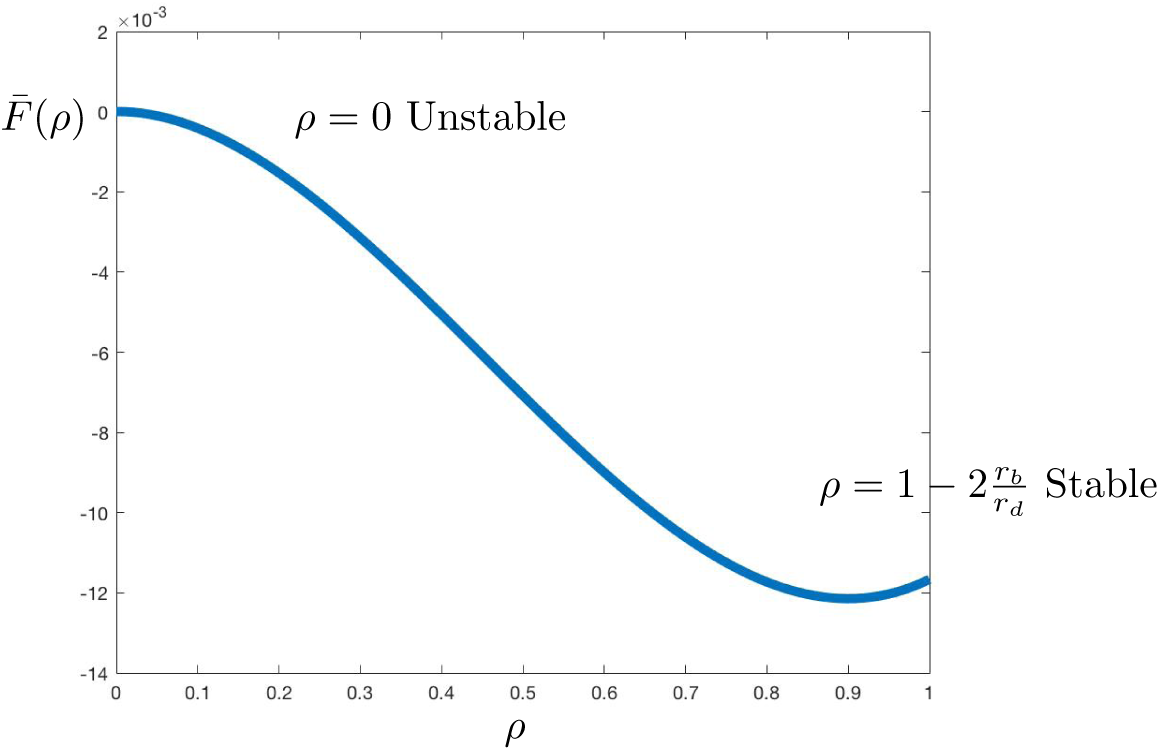
Plot of the potential of equation (5.10), namely 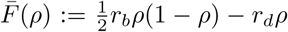 with *r*_*b*_ = 0.1 and *r*_*d*_ = 0.02.

### 5.3 Impact of heterogeneity on a spatio-temporal GoG model

In the previous section, and as a first step towards the investigation of the GoG model (5.6), spatial dependence was ignored and in consequence we investigated an ODE model, where only the dynamics of resting and switching between the two species were taken into account. Nonetheless, the main goal of the current section is to investigate what is the impact, if any, of the diffusion component on the stability analysis of the GoG model. Namely, in the current section we consider the full model (5.6) where now a randomization parameter is introduced both in the diffusion and the reaction terms. To do this, we follow again the averaging approach introduced in section 4. To guarantee the well posedness of the system, we again consider the functional approximation of the white noise introduced in (2.4).

Thus we focus on the investigation of the following

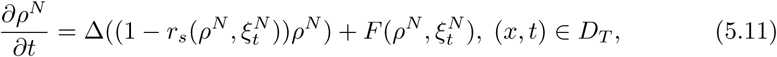

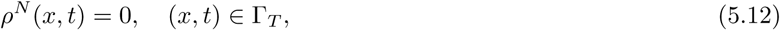

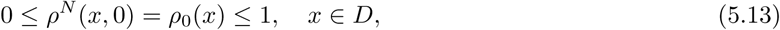

for a bounded and smooth *D* ⊂ ℝ^3^, where 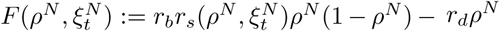. Here *T* > 0 again stands for the maximum existence time of solution *ρ*^*N*^. Note that the Dirichlet type boundary condition (5.12) means that there are no cancerous cells on the boundary of the domain under investigation *D*, which is a plausible assumption. Using [5, Theorem 9 in Chapter 7] we immediately obtain existence, uniqueness and positivity of a global-in-time solution, since also the drift term 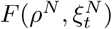 is bounded, for the random partial differential equation (RPDE) problem (5.11)-(5.13).

By virtue of an energy approach or via the maximum principle, we derive the positivity and boundeness of the solution of the limiting problem, while its uniqueness is derived using Gronwall’s lemma, see also,^21^

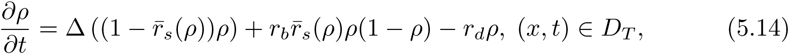

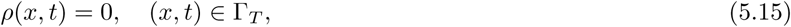

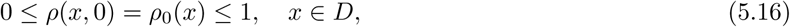

where 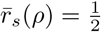.

Denote *H* = *L*^2^ (*D*) and ℋ_*T*_ = *C*(0, *T*; *H*) then making use of the novel averaging principle developed in section 4 we can prove the following averaging result, which is inspired by,^3^

#### Theorem 5.1

*For every T* > 0 *the unique solution ρ*^*N*^ *of problem* (5.11)*-*(5.13) *converges as N* → ∞ *into ℋ*_*T*_ *to the unique solution ρ of the average problem* (5.14)*-*(5.16).

**Proof.** First note that the term 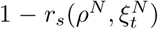 is bounded due the definition of 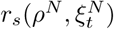 given by (5.4) and hence both the diffusion coefficient and the reaction term in problem (5.11)-(5.12) are also bounded. Thus via parabolic regularity we derive the a priori estimate

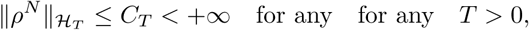

therefore we can find a subsequence denoted again by (*ρ*^*N*^) without any confusion and a function *ψ* ∈ ℋ_*T*_ such that

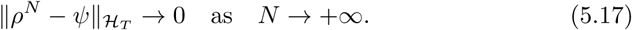

Considering the weak formulation of (5.11)-(5.13) we have

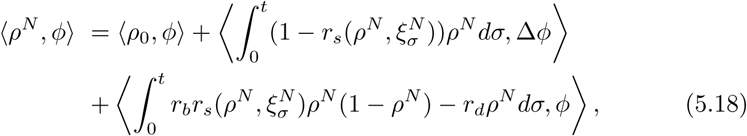

for any test function 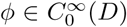, where ⟨·, ·⟩ stands for the inner product in *H* = *L*^2^ (*D*).

Using some algebraic manipulations then (5.18) infers

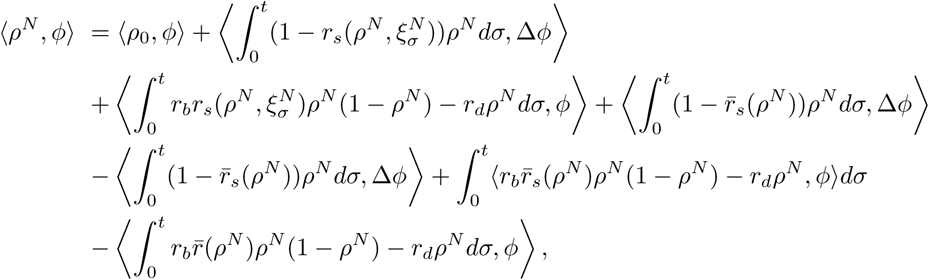

and after rearranging the preceding relation is reduced to

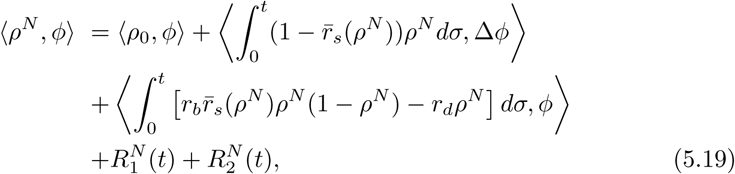

where

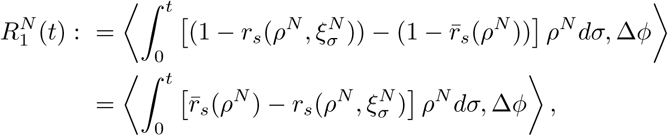

and

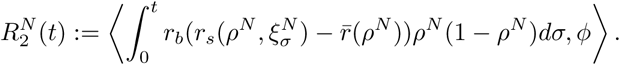

Considering the term 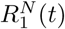 we derive the following estimate:

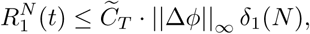

where 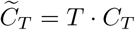 and

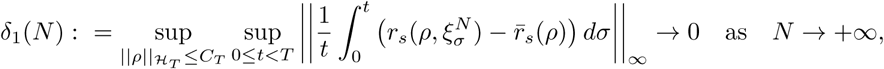

for any *T* > 0 by virtue of Lemma 5.1. Therefore for given *T* > 0

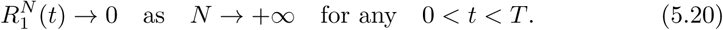

In a similar manner we have

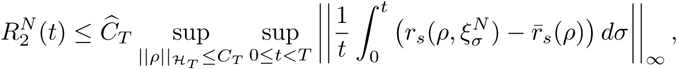

for any *T* > 0 where *Ĉ*_*T*_ := *r*_*b*_ *TC*_*T*_ (1 + *C*_*T*_ )‖*ϕ*‖_∞_ and using again Lemma 5.1 we infer

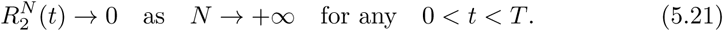

Letting now *N* → +∞ into (5.19) and using (5.17), (5.20) and (5.21) we derive

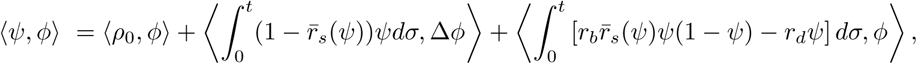

i.e. *ψ* is a weak solution of (5.14)-(5.16), and since this problem has a unique solution we finally infer that *ψ* = *ρ*. This completes the proof of the Theorem 5.1.

By the linear stability of problem (5.14)-(5.16), since the diffusion coefficient is small (c.f.^19^), we again obtain that 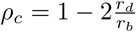 is stable whilst the trivial steadystate *ρ*_*nc*_ = 0 is unstable. Therefore, the diffusion has no effect on the stability of the spatial homogeneous steady-sates and again the cancerous population survives.

## 6. Discussion

In the current paper we have introduced a novel average method to deal with the study of the dynamics of a class of RODEs and RPDEs since classical averaging methods fail to treat this kind of problems. To elucidate the relevance of our theoretical results, we apply this method into an important biomedical problem, i.e. the assessment of intratumoral heterogeneity impact on tumor dynamics. In particular, we consider the development of gliomas according to a well-known Go or Grow (GoG) model, where intratumoral heterogeneity is modelled as a stochastic process. The deterministic version of the considered GoG model has been shown that there exhibits an emerging Allee effect (bistability). On the other hand, for the novel stochastic version of GoG model we demonstrate that the introduction of white noise, as a model of intratumoral heterogeneity, leads to a monostable tumor growth. The latter entails the disappearance of the Allee effect, and thus we conclude that the extinction is impossible under parametric variations. Consequently, our results suggest that the existence of heterogeneity worsens the prognosis of tumor growth, which is actually in accordance with the clinical experience and literature. However, the assumption of white noise is the worst possible scenario related to tumor heterogeneity, and therefore other noise distributions should be analyzed such as Gaussian noise. Moreover, there have been a plethora of studies trying to quantify intratumoral heterogeneity, see,^1, 20, 22, 24^ nevertheless our method is able to include the existing literature and analyze the impact of data-driven heterogeneity distribution in more realistic tumor models. Furthermore, our method can be also implemented to investigate the long-time dynamics of the full GoG system (5.1)-(5.2), which will be the subject of a forthcoming work.

## Acknowledgment

Part of the current work was inspired and initiated when NK was visiting the Center of Interdisciplinary Research (ZIF) and Helmholtz Centre for Infection Research. He would like to express his gratitude for the warm hospitality of both institutes and the financial support from ZIF. HH is supported by Mic2Mode-I2T (01ZX1710B) and HH is supported by SYSIMIT (01ZX1308D) and MulticellML (01ZX1707C) by the Federal Ministry of Education and Research (BMBF) and by the SYSMIFTA (031L0085B) of the ERACOSYSMED initiative. JCLA and HH gratefully acknowledges the funding support of the Helmholtz Association of German Research Centers - Initiative and Networking Fund for the project on Reduced Complexity Models (ZT-I-0010)

## References

1. L. Alic, W. J. Niessen and J. F. Veenland, Quantification of heterogeneity as a biomarker in tumor imaging: A systematic review, PLoS ONE 9 (2014) 1–15.

2. K. Böttger, H. Hatzikirou, A. Voss-Böhme, E. A. Cavalcanti-Adam, M. A. Herrero and A. Deutsch, An Emerging Allee Effect Is Critical for Tumor Initiation and Persistence, PLOS Computational Biology 11 (2015) e1004366.

3. J. Duan and W. Wang, Effective Dynamics of Stochastic Partial Differential Equations(Elsevier, 2014).

4. M. I. Freidlin and A. D. Wentzell, Random perturbations, in Random perturbations of dynamical systems(Springer, 1998), pp. 15–43.

5. A. Friedman, Partial Differential Equations of Parabolic Type (Prentice-Hall, Inc., Englewood Cliffs, N.J., 1964).

6. J. C. L. Alfonso, A. Köhn-Luque, T. Stylianopoulos, F. Feuerhake, A. Deutsch and H. Hatzikirou, Why one-size-fits-all vaso-modulatory interventions fail to control glioma invasion: in silico insights, Scientific Reports 6 (2016) 37283.

7. I. M. Gelfand and G. E. Shilov, Generalized functions, Vol. 4: applications of harmonic analysis(Academic Press, 1964).

8. J. C. L. Alfonso K. Talkenberger, M. Seifert, B. Klink, A. Hawkins-Daarud, K. R. Swanson, H. Hatzikirou, A. Deutsch The biology and mathematical modelling of glioma invasion: a review, J. R. Soc. Interface 14(136) (2017) 1–20.

9. M. L. Goodenberger and R. B. Jenkins, Genetics of adult glioma, Cancer genetics 205 (2012) 613–621.

10. J. Hale and S. Verduyn Lunel, Averaging in infinite dimensions, Journal of Integral Equations and Applications 2 (1990) 463–494.

11. H. Hatzikirou, J. C. L. Alfonso, S. Mühle, C. Stern, S. Weiss and M. Meyer-Hermann, Cancer therapeutic potential of combinatorial immuno- and vaso-modulatory interventions, J. R. Soc. Interface 12 (2015) 20150439.

12. X. Han and P. E. Kloeden, Random Ordinary Differential Equations and Their Numerical Solution, Probability Theory and Stochastic Modelling 85 (Springer, 2017).

13. H. Hatzikirou, D. Basanta, M. Simon, K. Schaller and A. Deutsch, go or grow: the key to the emergence of invasion in tumour progression?, Mathematical medicine and biology: a journal of the IMA 29 (2012) 49–65.

14. D. Henry, Geometric Theory of Semilinear Parabolic Equations, Lecture Notes in Mathematics (Springer Berlin Heidelberg, 2006).

15. W. S. Ikeda N., Stochastic Differential Equations and Diffusion Processes(North-Holland Publishing Co., Amsterdam, 1981).

16. D. N. Louis, H. Ohgaki, O. D. Wiestler, W. K. Cavenee, P. C. Burger, A. Jouvet, B. W. Scheithauer and P. Kleihues, The 2007 who classification of tumours of the central nervous system, Acta neuropathologica 114 (2007) 97–109.

17. N. McGranahan and C. Swanton, Biological and therapeutic impact of intratumor heterogeneity in cancer evolution, Cancer Cell 27 (2015) 15–26.

18. C. E. Meacham and S. J. Morrison, Tumour heterogeneity and cancer cell plasticity, Nature 501 (2013) 328–337.

19. J. Murray, Mathematical Biology Vol II:Spatial Models and Biomedical Applications(Springer, 2003).

20. L. Oesper, G. Satas and B. J. Raphael, Quantifying tumor heterogeneity in whole-genome and whole-exome sequencing data, Bioinformatics 30 (2014) 3532–3540.

21. P. Quittner and S. Ph., Superlinear parabolic problems. Blow-up, global existence & steady states, Birkhauser Adv. Texts Basler Lehrbücher (Birkhäuser, 2007).

22. E. M. Rutter, H. T. Banks and K. B. Flores, Estimating intratumoral heterogeneity from spatiotemporal data, Journal of Mathematical Biology.

23. N. Sanai, A. Alvarez-Buylla and M. S. Berger, Neural stem cells and the origin of gliomas, New England Journal of Medicine 353 (2005) 811–822.

24. A. Sottoriva, I. Spiteri, S. G. M. Piccirillo, A. Touloumis, V. P. Collins, J. C. Marioni, C. Curtis, C. Watts and S. Tavaré, Intratumor heterogeneity in human glioblastoma reflects cancer evolutionary dynamics., Proceedings of the National Academy of Sciences of the United States of America 110.

25. M. Tektonidis, H. Hatzikirou, A. Chauvière, M. Simon, K. Schaller and A. Deutsch, Identification of intrinsic in vitro cellular mechanisms for glioma invasion., Journal of theoretical biology 287 (2011) 131–47.

